# Population Pharmacokinetics and Target Attainment Analysis of Vancomycin after Intermittent Dosing in Adults with Cystic Fibrosis

**DOI:** 10.1101/2023.07.28.551040

**Authors:** Venkata K. Yellepeddi, Bryn Lindley, Emi Radetich, Shaun Kumar, Zubin Bhakta, Laurie Leclair, Madison Parrot, David C. Young

**Author notes:** **Corresponding Author** David C. Young, Professor (Clinical), Department of Pharmacotherapy College of Pharmacy, University of Utah, L.S. Skaggs Pharmacy Institute, Room 4916, Salt Lake City, Utah – 84112.

## Abstract

Vancomycin is the first-line agent to treat pulmonary infections caused by methicillin-resistant *Staphylococcus aureus* (MRSA) in people with cystic fibrosis (PwCF). However, there is no consensus on vancomycin dosing in this population among health institutions, and there is large variability in dosing regimens across the United States. In this study, we characterized the pharmacokinetics (PK) of vancomycin in PwCF using a population PK approach. The clinical PK data to develop the population PK model was obtained from vancomycin therapeutic monitoring data from PwCF undergoing treatment for infections due to MRSA. The population PK model was then used to perform comprehensive Monte Carlo simulations to evaluate the probability of target attainment (PTA) of 12 different dosing scenarios. The area under the curve to minimum inhibitory concentration ratio (AUC/MIC) ≥ 400 mg*h/L was used as a target for PTA analysis. A total of 181 vancomycin plasma concentrations were included in the analysis. A one-compartment model with first-order elimination best described the data. Weight significantly influenced the vancomycin PK (*p* < 0.05). In the final model, clearance was estimated as 5.52 L/h/70 kg, and the volume of distribution was 31.5 L/70 kg. The PTA analysis showed that at lower MIC levels (MIC = 1), doses greater than and equal to 1000 mg every 8 hours and 1250 mg every 12 hours resulted in >90% PTA. The PTA results from this study may potentially inform the design of vancomycin dosing regimens to treat pulmonary infections due to MRSA in PwCF.

## INTRODUCTION

In PwCF, a leading cause of pulmonary exacerbations is due to MRSA infections (1, 2). In the United States, the prevalence of MRSA rapidly increased from 2% in 2001 to ∼26.1% in 2016, with a current prevalence of 18% in 2021 (3, 4). Unfortunately, PwCF infected by MRSA have a 27% higher risk of death and 6.2-year shorter life than their non-infected counterparts (5). The American Thoracic Society Guidelines on antibiotic management of lung infections in CF recommend intravenous (IV) vancomycin as the first-line treatment of MRSA in PwCF (2). However, there are no appropriate vancomycin dosing guidelines for MRSA in PwCF, and the dosing in this population widely varies among different health institutions, from 20 to 200 mg/kg/day IV every 6 to 12 hours in pediatric patients and 500 to 2000 mg IV every 6 to 12 hours in adult patients (6). The need to optimize the vancomycin dosing regimens and monitoring strategies in PwCF has been highlighted in the National Practice Survey conducted at various CF centers across the United States (7).

Vancomycin is a tricyclic glycopeptide antibiotic widely used to treat bacterial infections caused by gram-positive bacteria, including MRSA (8). After IV administration, vancomycin is eliminated primarily via the renal route, with >80% recovered unchanged in urine within 24 h (9). In patients with normal renal function, vancomycin plasma half-life is approximately 4 to 6 hours with a clearance of 0.058 L/kg/h and volume of distribution (V_d_) ranging from 0.4 - 1 L/kg (10). The protein binding of vancomycin ranges from 10% to 50% (10). Nephrotoxicity and ototoxicity are the major dose-limiting toxicities of vancomycin that warrant careful consideration while dosing this drug (8). Despite the variability in dosing, guidelines recommend therapeutic drug monitoring of vancomycin with a goal of trough serum concentrations between 15 to 20 mg/L (6). However, data from experimental and clinical studies have selected the AUC/MIC as the best parameter to predict the efficacy of vancomycin with a target consensus of an AUC/MIC ratio of ≥400 mg*h/L (11-13). The reported data also suggested that the risk of vancomycin-associated nephrotoxicity is significantly increased at AUC_0-24_ > 600-700 mg*h/L (14, 15).

The pharmacokinetics (PK) of antibiotics used in PwCF has been reported to differ from people without CF due to physiological and disease factors such as chronic inflammation, impaired intestinal bicarbonate secretion, decreased plasma protein binding, increased renal clearance, and increased glomerular filtration rate (4). Therefore, it is very important to characterize the pharmacokinetics of antibiotics in this population and use that information to design a dosing regimen to maximize therapeutic efficacy and minimize toxicity. To date, only two studies have reported the pharmacokinetics of vancomycin in adult PwCF (16, 17). In 1996, Pleasants et al. reported vancomycin PK in PwCF after a single IV dose in 10 patients (16). More recently, we reported the PK of vancomycin in PwCF using retrospective PK data for vancomycin after intermittent dosing in PwCF with MRSA pulmonary infection (17). However, there are no reports in the literature that characterized the population PK and assessed the target attainment analysis of IV intermittent-dosed vancomycin in adult PwCF.

There is a significant unmet medical need to evaluate the appropriateness of the current vancomycin dosing regimens to treat MRSA infections in PwCF. To address this unmet medical need, for the first time, we are reporting population pharmacokinetics and the probability of target attainment (PTA) of vancomycin in adult PwCF. In this study, we developed a population PK model and evaluated the covariates influencing the PK of vancomycin in adult PwCF at the University of Utah Health Adult Cystic Fibrosis Center. The PTA analysis at various vancomycin dosing regimens was performed using the population PK model using Monte Carlo simulation. The target for PTA included an efficacy target of AUC_0-24_/MIC ≥400 mg*h/L and a safety target of AUC_0-24_ < 650 mg*h/L.

## MATERIALS AND METHODS

### Study population

The data were collected retrospectively from adult PwCF admitted to the University of Utah Hospital between May 2014 and August 2020 for pulmonary exacerbations (PEx). Individuals were included if at least two vancomycin serum concentrations were recorded. Patient demographics, including sex, age, weight, height, serum creatinine, and use of concomitant inhaled antibiotics such as tobramycin and aztreonam were collected from the electronic data warehouse. Creatinine clearance was calculated using the Cockcroft-Gault formula (18). Patients were excluded if they were pregnant as determined by elevated serum human chorionic gonadotropin levels. Patients were also excluded if they received IV vancomycin for less than 24 hours. This study was reviewed and approved by the University of Utah Institutional Review Board (IRB#00136598).

### Dosing and sampling schedule

Vancomycin doses 500, 750, 1000, 1250, 1500, 1750, and 2000 mg were administered by IV infusion at a standard rate of 500 mg/hour at 8-or 12-hour intervals. The blood samples to analyze peak vancomycin levels were collected at least 1 hour after infusion completion and blood samples to analyze trough vancomycin levels were drawn 1 hour prior to the next dose. PwCF admitted for PEx received care for 10 to 14 days in the inpatient setting at the University of Utah Hospital. Because patients are often co-infected with *Pseudomonas aeruginosa*, inhaled antibiotics tobramycin and aztreonam were included in the covariate analysis and were administered to patients by nebulization at doses: 300 mg every 12 hours for tobramycin and 75 mg every 8 hours for aztreonam. Inhaled vancomycin is held during admission for PEx, so was not included in the analysis.

### Vancomycin assay

Serum vancomycin concentrations were measured using a MULTIGENT vancomycin assay on ARCHITECT cSystem by Abbott Laboratories, IL, USA. The MULTIGENT assay is a homogenous particle-enhanced turbidimetric inhibition immunoassay (PETNIA). The assay validated for clinical use and the lower limit of quantitation (LLOQ) for vancomycin assay was 1.1 mg/L. The assay was linear within the vancomycin concentration range of 1.1 to 100 mg/L. The intra- and interday precision for vancomycin assay was 1.38% and 1.52% respectively. The percentage recovery of vancomycin across a concentration range of 2.5 to 75 mg/L ranged from 99.29% to 105.2%.

### Pharmacokinetic model development

The population PK analysis for vancomycin plasma concentrations was performed using a nonlinear mixed-effect modeling approach using software NONMEM^®^ 7.5, ICON plc., MD, USA interfaced with Perl-speaks-NONMEM (PsN^®^) version 5.3.0 (19), and Finch Studio™ NONMEM^®^ workbench, Enhanced Pharmacodynamics LLC, NY, USA (20). The pre- and post-processing of the datasets and development of graphics was performed using Tidyverse package in R software (21) with R Studio^®^ interface (22). First order conditional estimation method with interaction was used to estimate the population values of the PK parameters, interindividual variability (IIV), and residual unexplained variability (RUV). Prediction corrected visual predictive checks (VPCs) were performed with (PsN^®^) version 5.3.0.

#### (i) Structural and error models

The base structural model was evaluated using different compartment models (e.g., one-compartment versus two-compartment) with first-order elimination. Diagnostic plots were used to visually inspect the model’s fit, including observed versus population predicted vancomycin concentrations and observed versus individual predicted vancomycin concentrations. Residuals and conditional weighted residuals were also plotted versus time or population predicted vancomycin concentrations. Models were further compared by assessing the precision of the parameter estimates, measures of variability, and the objective function value (OFV).

Model variability and random effects were classified as one of two types of error: interindividual variability (IIV) and residual unexplained variability (RUV). The IIV is the variability inherent between different patients and was assumed to be log-normally distributed according to an exponential equation of the form:

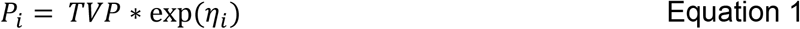

Where (P_i_) is the individual PK parameter, (TVP) is the typical population mean of the PK parameter, and (η_i_) is the proportional difference between the *i*^th^ subject’s parameters estimate and the typical population mean. Eta is assumed to be normally distributed with a mean of 0 and a variance of ω^2^ (23).

RUV reflects the difference between the model prediction for the individual and the measured observation. This includes the error in the assay, errors in drug dose, errors in the time of measurement etc. (24). During model development, RUV was evaluated using additive, proportional, and combined error models. Proportional error model resulted in the greatest improvement in the OFV. The equation for the proportional error model was:

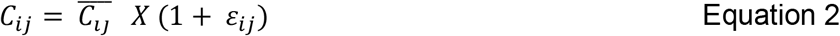

Where (C_ij_) is the individual observed plasma concentration at time j, 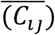 is the individual predicted plasma concentration at time j, and (ε_*ij*_) is the proportional error, which is assumed to be normally distributed with a mean of 0 and a variance of σ^2^.

The area under the plasma concentration versus time curve over a day (AUC_24_) for vancomycin was calculated for each patient using the following equation:

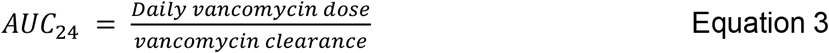

#### (ii) Covariate analysis

The covariates evaluated in the analysis include age, weight, creatinine clearance, and concomitant use of inhaled antimicrobials. The inhaled antimicrobials included tobramycin alone, aztreonam alone, or both tobramycin and aztreonam. Race was not included in the covariate analysis as all patients in the dataset were white. Sex was not included as there was no difference in exploratory analysis plots between male and female patients. Further testing was performed by evaluating potential covariates using stepwise forward addition and then stepwise backward elimination procedures. A reduction in the OFV of more than 3.84 (p<0.05) was required to retain covariates in the forward addition step. In the backward elimination step, covariates were retained if they resulted in a reduction in the OFV of more than 10.83 (p<0.001). For continuous covariates age, weight, and creatinine clearance power model was considered:

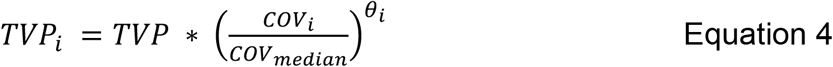

Where TVP_i_ represents the individual PK parameter, (TVP) represents the typical value of the parameters, and (θ_cov_) represents the scale factor for the individual covariate (COV_i_) and the median covariate value COV _median_.

The model for a categorical covariate concomitant inhaled antibiotics was expressed using a proportional model:

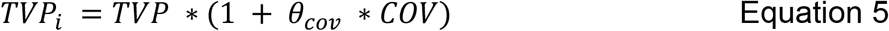

Where TVP represents the typical value of the parameters, and (*θ*_*cov*_) represents the fractional change in the typical parameter with the covariate (COV), where COV has at least two or more levels.

#### (iii) Model evaluation

The likelihood ratio test (−2LLR) was used to calculate the difference between the nested models, difference in objective function values (ΔOFV), and between models with a decrease ≥ 3.84 which is statistically significant at an α = 0.05. Nonnested models were evaluated by using Akaike information criterion (AIC). Final selection of models was also based on goodness-of-fit plots along with the reliability of the parameters estimates, such as reduced variability between subjects and residual errors.

A prediction corrected VPC was performed to evaluate the final model. The original dataset and final model parameter estimates were used to simulate 1,000 individual datasets. The observed concentrations and the median (50^th^ percentile) and 5^th^ and 95^th^ percentiles were plotted along with the simulated 95% confidence intervals of the median and 5^th^ and 95^th^ percentiles. The final model is considered appropriate if the observed data falls within the simulated 95% confidence intervals of the 5^th^, 50^th^, and 95^th^ percentiles of the simulated concentrations.

### Model simulations

Based on the final population PK model for vancomycin, Monte Carlo simulations were performed using NONMEM to evaluate the PTA for different dosing regimens of vancomycin. The dosing regimens simulated are: 500, 750, 1000, 1250, 1500, and 2000 mg every 8 and 12 hours. The primary PK/PD target for vancomycin included AUC/MIC ≥ 400. In total, 12 different dosing regimens were evaluated. The MIC values covered a range of 0.125 to 8 mg/L based on the reported range of vancomycin MIC’s for MRSA (25). PTA was evaluated by determining the percentage of the simulated patient population in which the specified PK/PD target (i.e., AUC/MIC ≥ 400 mg*h/L) was achieved. The PTA was also evaluated by including a toxicity threshold of AUC_0-24_ < 650 mg*h/L to the PK/PD target.

## RESULTS

### Patient characteristics

A total of 181 plasma samples were obtained from 19 patients. All concentrations were quantified above the LLOQ of 1.1 mg/L. Demographic data collected in this study is shown in Table 1. There were 5 males and 19 females with a median age (range) of 27 (19-60) years. All the patients identified themselves as white. There was a total of 110 vancomycin doses administered among the 19 included patients. All patients received at least two doses of vancomycin. The percentage of the total number of doses administered was highest for 750 mg (37.2%) and lowest for 1750 mg. The table with percentage of total number doses stratified based on individual dose amount is provided as a table in supplementary datafile S1. The mean of vancomycin levels was 21.4 ± 9.6 mg/L and the median (range) of vancomycin levels was 19.8 (6.6 – 61.4) mg/L.

**Table 1.**
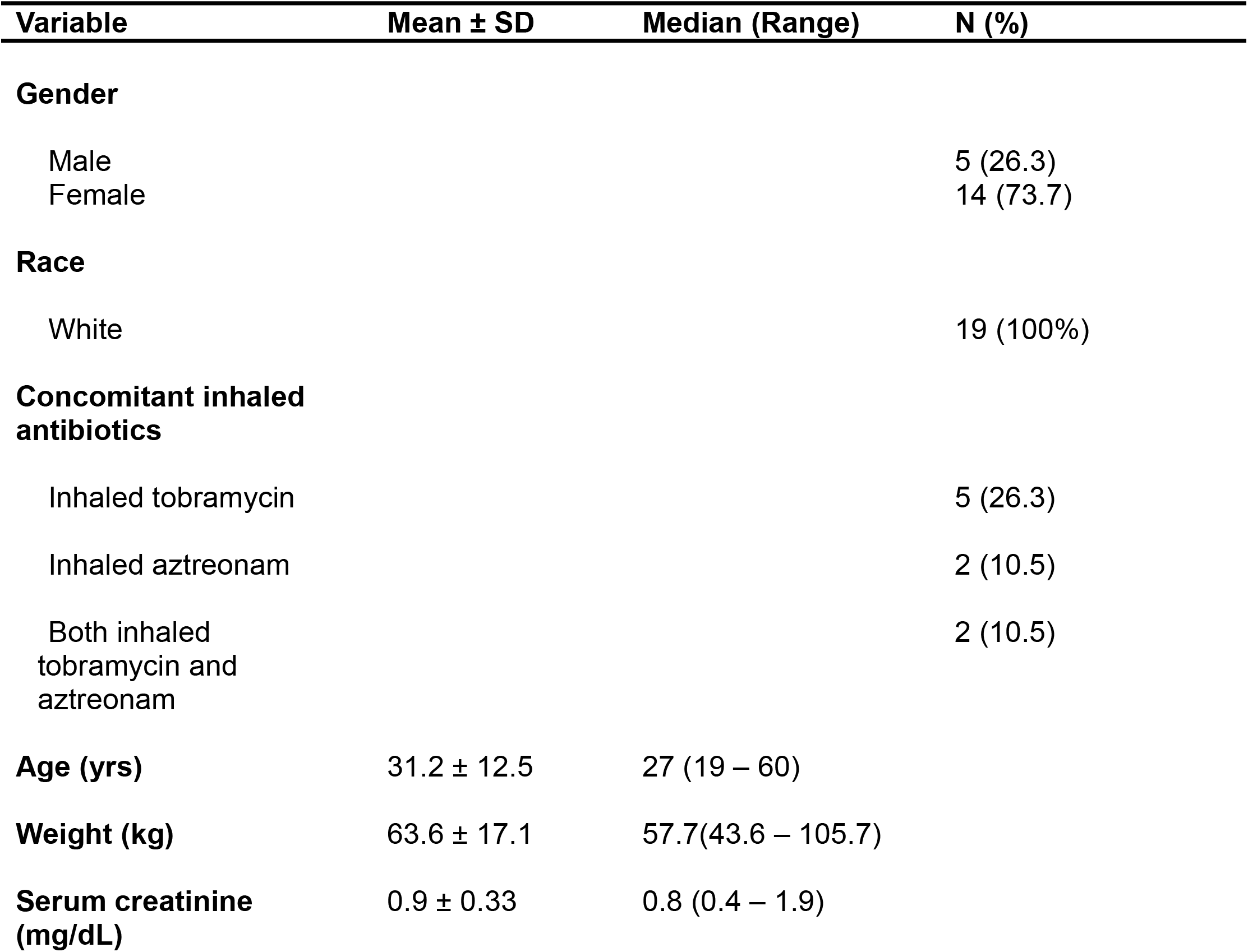
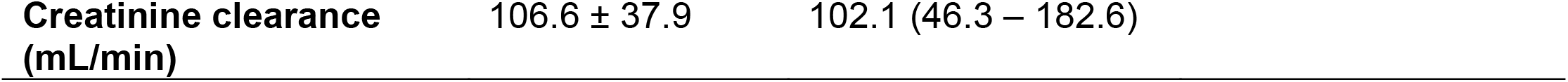
Demographic characteristics in adult PwCF who received vancomycin for the treatment of a pulmonary exacerbation.

| Variable | Mean ± SD | Median (Range) | N (%) |
| --- | --- | --- | --- |
| **Gender** |  |  |  |
| Male |  |  | 5 (26.3) |
| Female |  |  | 14 (73.7) |
| **Race** |  |  |  |
| White |  |  | 19 (100%) |
| **Concomitant inhaled antibiotics** |  |  |  |
| Inhaled tobramycin |  |  | 5 (26.3) |
| Inhaled aztreonam |  |  | 2 (10.5) |
| Both inhaled tobramycin and aztreonam |  |  | 2 (10.5) |
| **Age (yrs)** | 31.2 ± 12.5 | 27 (19 – 60) | |
| **Weight (kg)** | 63.6 ± 17.1 | 57.7(43.6 – 105.7) | |
| **Serum creatinine (mg/dL)** | 0.9 ± 0.33 | 0.8 (0.4 – 1.9) | |

|  |  |  |
| --- | --- | --- |
| **Creatinine clearance**<br>**(mL/min)** | 106.6 ± 37.9 | 102.1 (46.3 – 182.6) |

### Population Pharmacokinetic Models

One-compartment and two-compartment structural models with first-order elimination were assessed with additive, proportional, and combined error models. Structural models also incorporate the rate and duration of the intravenous infusion for each patient. The base model was a one-compartment model with first-order elimination with a proportional error model, which was selected as the initial base model based on the OFV. IIV terms were placed on clearance and volume of distribution.

Results from the covariate analysis showed that weight (*p*<0.05) was identified as having significant influence on vancomycin PK. However, in multivariate analyses accounting for weight, adding creatinine clearance showed significant (*p*<0.05) influence but increased the %RSE on weight exponent by three times. Therefore, creatinine clearance was not included in the model. The summary of the sequential covariate model development is provided as a table in supplementary datafile S2. The final covariate model was chosen as it produced the most significant minimization of the OFV (Δ 8.27), reduced the IIV, and decreased the RUV. The parameter estimates derived from the final covariate model are shown in Table 2.

**Table 2.**
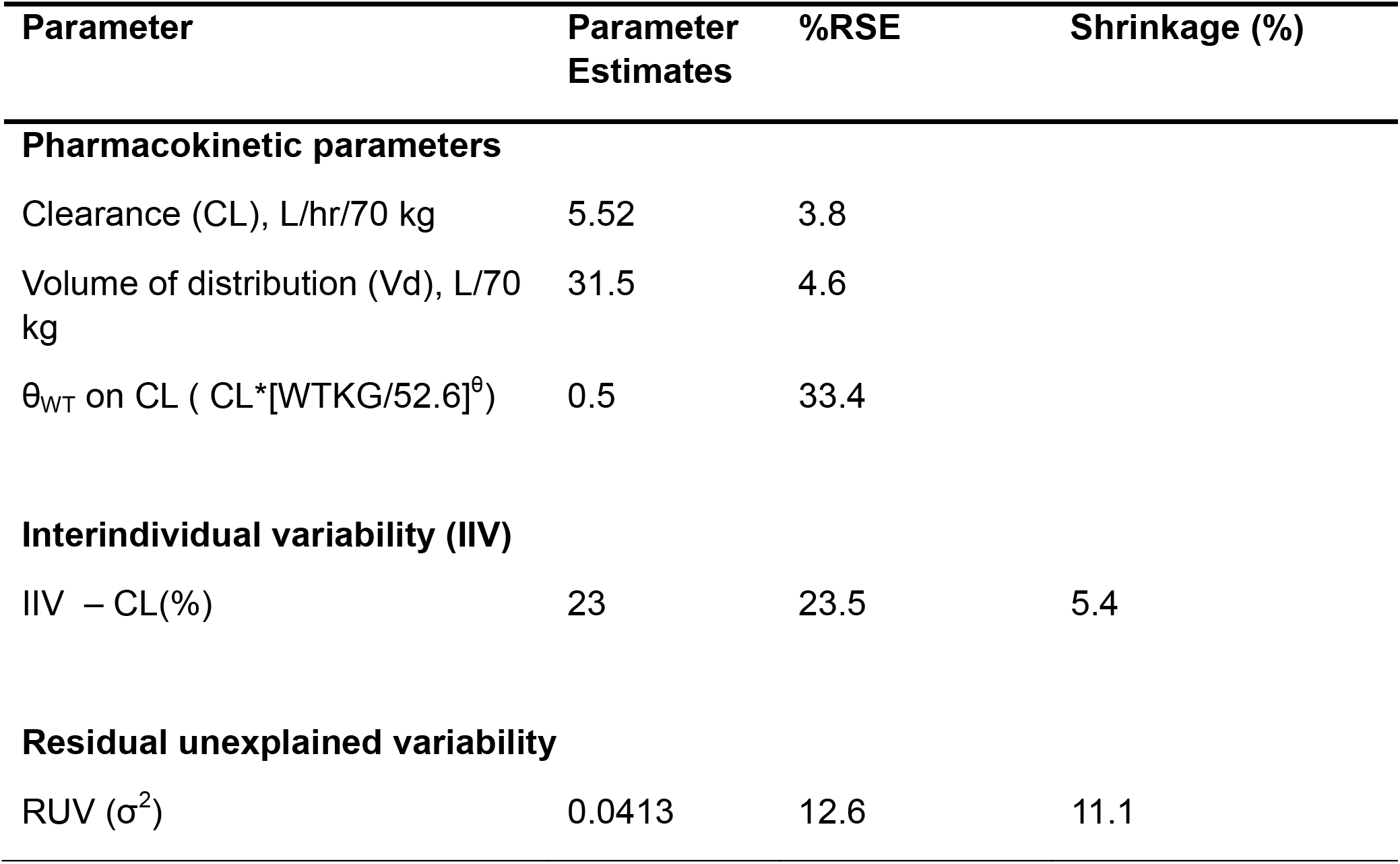
Final population PK model parameters estimates for vancomycin population PK model.

Diagnostic plots were generated for assessing model fit between observed vancomycin concentrations versus population predicted and individual predicted values (Figure 1). The conditional weighted residuals versus the population predicted vancomycin concentration plots were also examined (Figure 1). The observed plasma concentrations versus population-predicted and individual-predicted plasma concentrations evenly distributed around the line of unity, suggesting that there was no major bias in the stochastic models of structural model. Overall, visual inspection revealed that the final covariate model fit the data more tightly than the initial base model. The prediction corrected VPC plot, presented in Figure 2, showed that a high percentage of the observed concentrations were within predicted 95% confidence intervals of the simulated data, indicating reasonable model stability and agreement.

**Figure 1.**
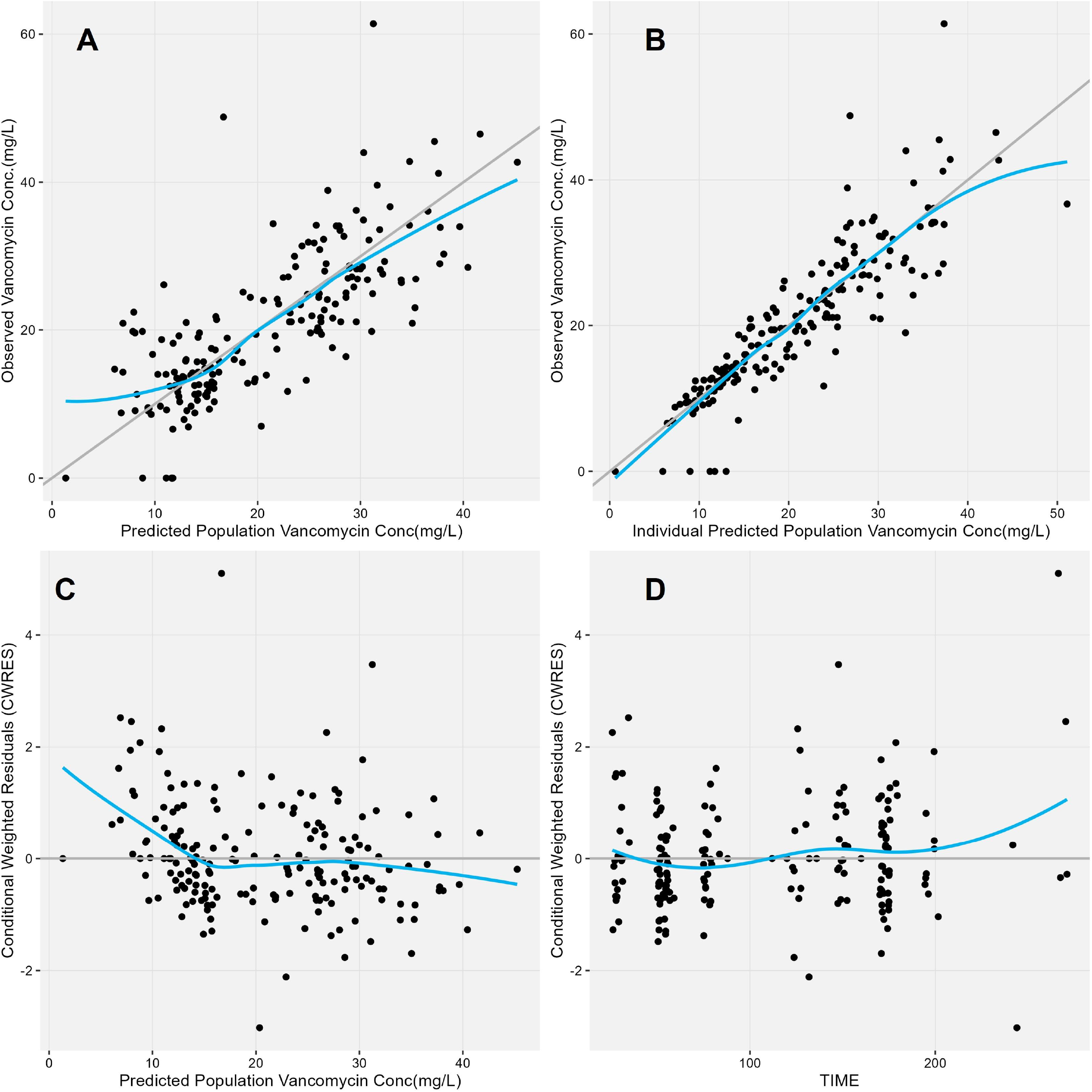
Diagnostic plots of the final-model (A) observed versus population predicted vancomycin concentrations, (B) observed versus individual predicted vancomycin concentrations (IPRED), (C) conditional weighted residuals (CWRES) versus population predicted vancomycin concentrations, and (D) CWRES versus the time after dose. Gray lines represent line of unity and blue lines represent loess smoothers.

**Figure 2.**
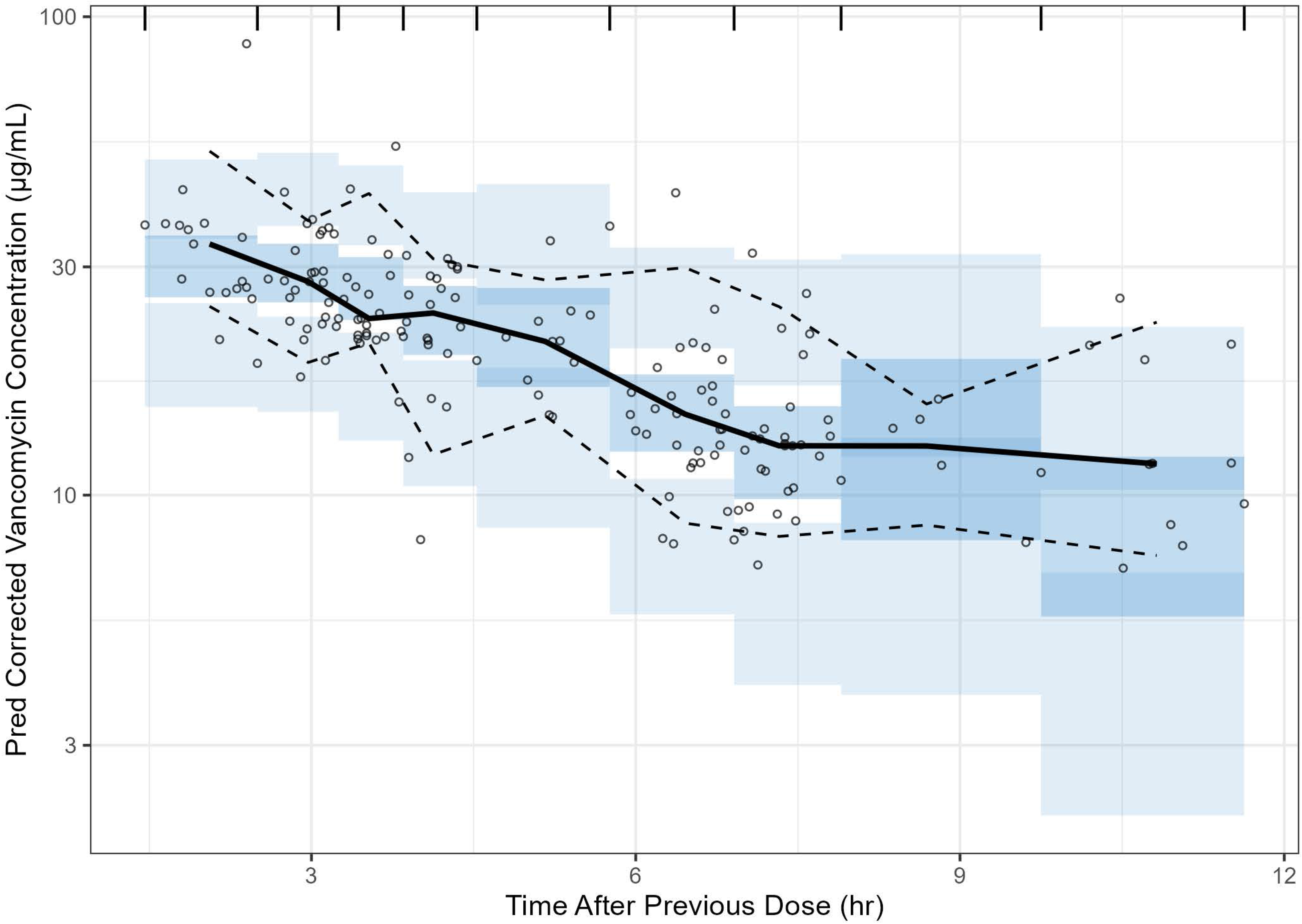
Prediction corrected visual predictive check (pcVPC) for final model. Black dots, observed values; solid black line, medial (50^th^ percentile); dashed black lines, 5^th^ and 95^th^ percentiles for the observed values, dark and light blue shaded regions, 95% confidence intervals for the 5^th^, 50^th^, and 95^th^ percentiles of the simulated data.

The probability of attaining bactericidal targets in plasma using various vancomycin dosing regimens across a wide range of MICs were evaluated with the target AUC/MIC ≥ 400. The PTA versus MIC profiles for different dosing schemes are provided in Figure 3. The simulations figure showed that at lower MIC levels (MIC = 1) doses greater than and equal to 1000 mg every 8 hours and 1250 mg every 12 hours resulted in >90% PTA. However, as the MIC values increases the % of PTA drastically decreased with lower dosing regimens. For instance, at MIC = 4, none of the dosing regimens achieved a PTA of >90%.

**Figure 3.**
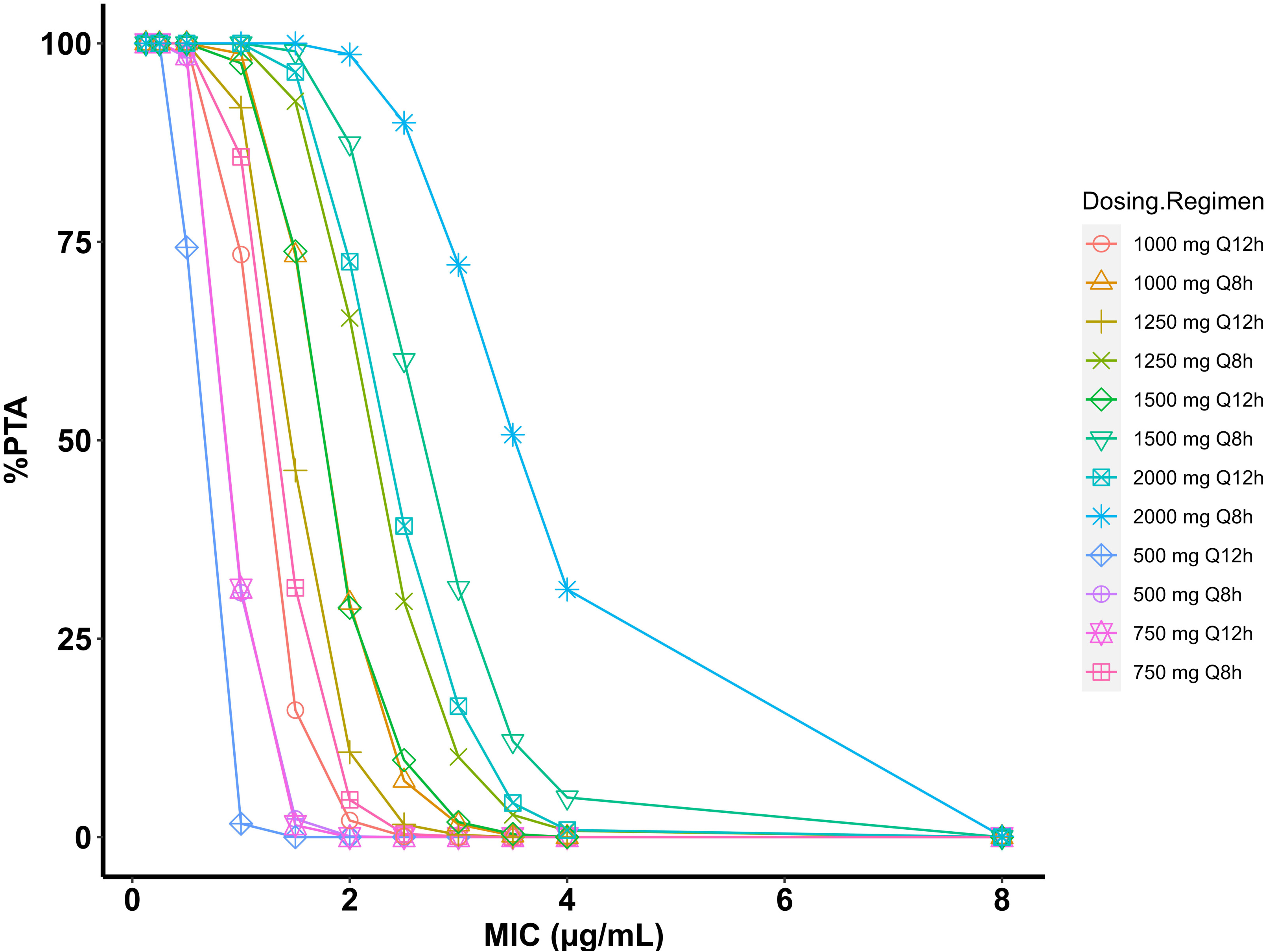
The probability of target attainments (PTA), achieving the target AUC_0-24_ ≥ 400, with different MIC values of vancomycin for MRSA using different vancomycin dosing regimens.

The PTA analysis was also performed by including a toxicity target of AUC_0-24_ > 650 mg*h/L. Table 3 shows the percentage of patients achieving the target with and without including the toxicity target at all dosing regimens simulated using the population PK (PoPPK) model. When toxicity target was included, the % PTA drastically reduced in dosing regimens greater than 750 mg for both 8- and 12-hour intervals. The highest %PTA of 57.4 % was achieved in 1000 mg q8h population. The violin plots showing the AUC_0-24_ values at various dosing regimens between 500-2000 mg for 8-hour and 12-hour intervals are provided as Figure 4 and Figure 5.

**Table 3.** %PTA of vancomycin following different dose regimens in simulated subjects^a^.

| Dosing Regimen<br>(mg) | Vancomycin Targets for PTA (%) |  |  |
| --- | --- | --- | --- |
| | $AUC_{0-24}/MIC \geq 400$<br>mg*h/L<sup>b</sup> | $AUC_{0-24} \geq 400$ and <<br>650 mg*h/L | $AUC_{0-24} > 650$<br>mg*h/L |
| 500 q8h | 30.8 | 28.5 | 2.3 |
| 750 q8h | 85.7 | 54.3 | 31.4 |
| 1000 q8h | 98.7 | 25.4 | 73.3 |
| 1250 q8h | 99.9 | 7.2 | 92.7 |
| 1500 q8h | 100 | 1 | 99 |
| 2000 q8h | 100 | 0 | 100 |
| 500 q12h | 1.7 | 1.7 | 0 |
| 750 q12h | 31.3 | 29.8 | 1.5 |
| 1000 q12h | 73.4 | 57.4 | 16 |
| 1250 q12h | 91.9 | 45.7 | 46.2 |
| 1500 q12h | 97.5 | 23.7 | 73.8 |
| 2000 q12h | 100 | 3.6 | 96.4 |

<sup>a</sup>The number of subjects included in the simulations was 1000 <sup>b</sup> For this calculation MIC was considered to be 1 µg/mL.

**Figure 4.**
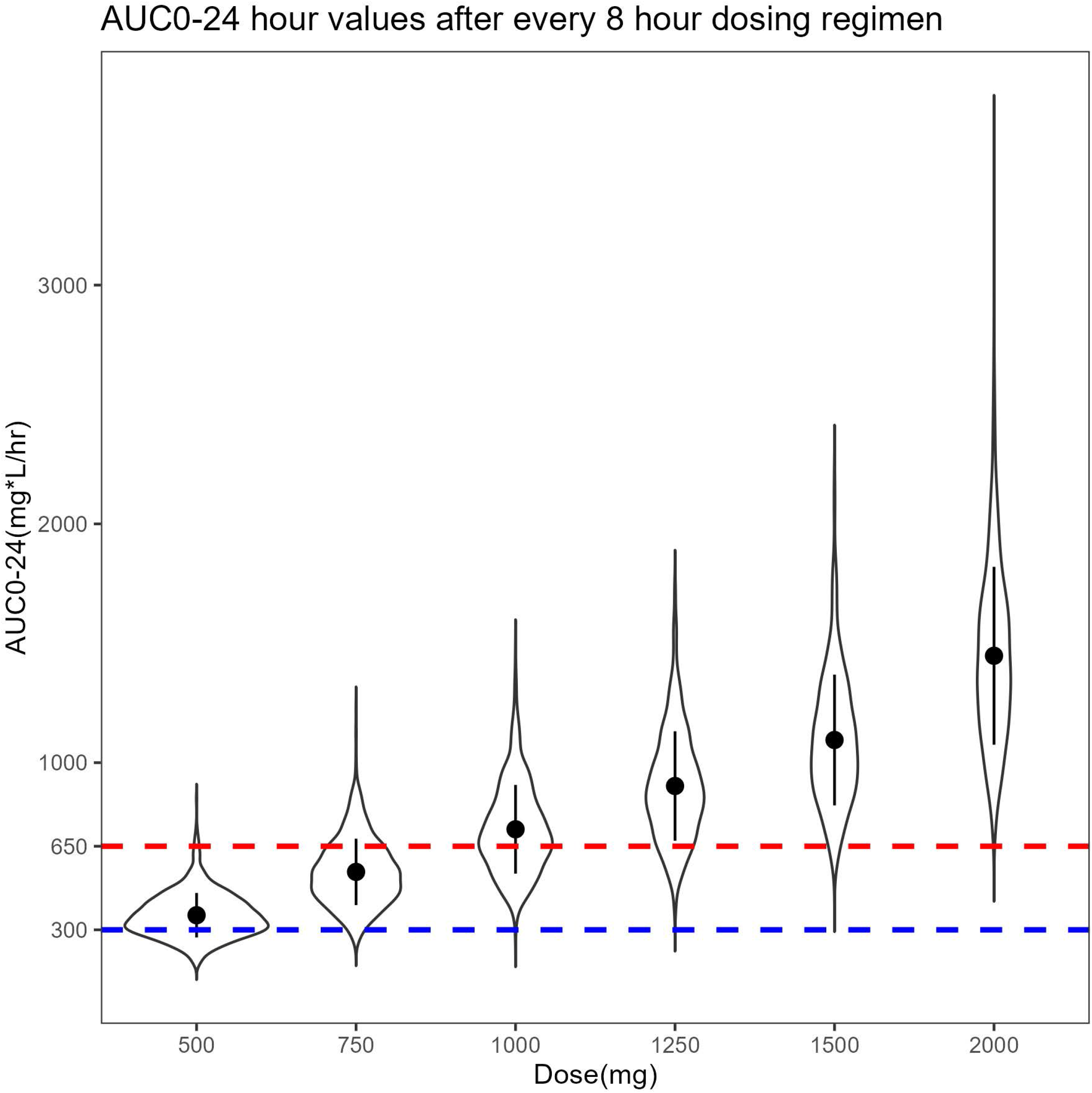
Violin plots of AUC_0-24_ versus various doses administered at every 8 hour interval. The black circle represents the median AUC_0-24_ and the black line represent the interquartile range. The white shaded area illustrate the kernel probability density.

**Figure 5.**
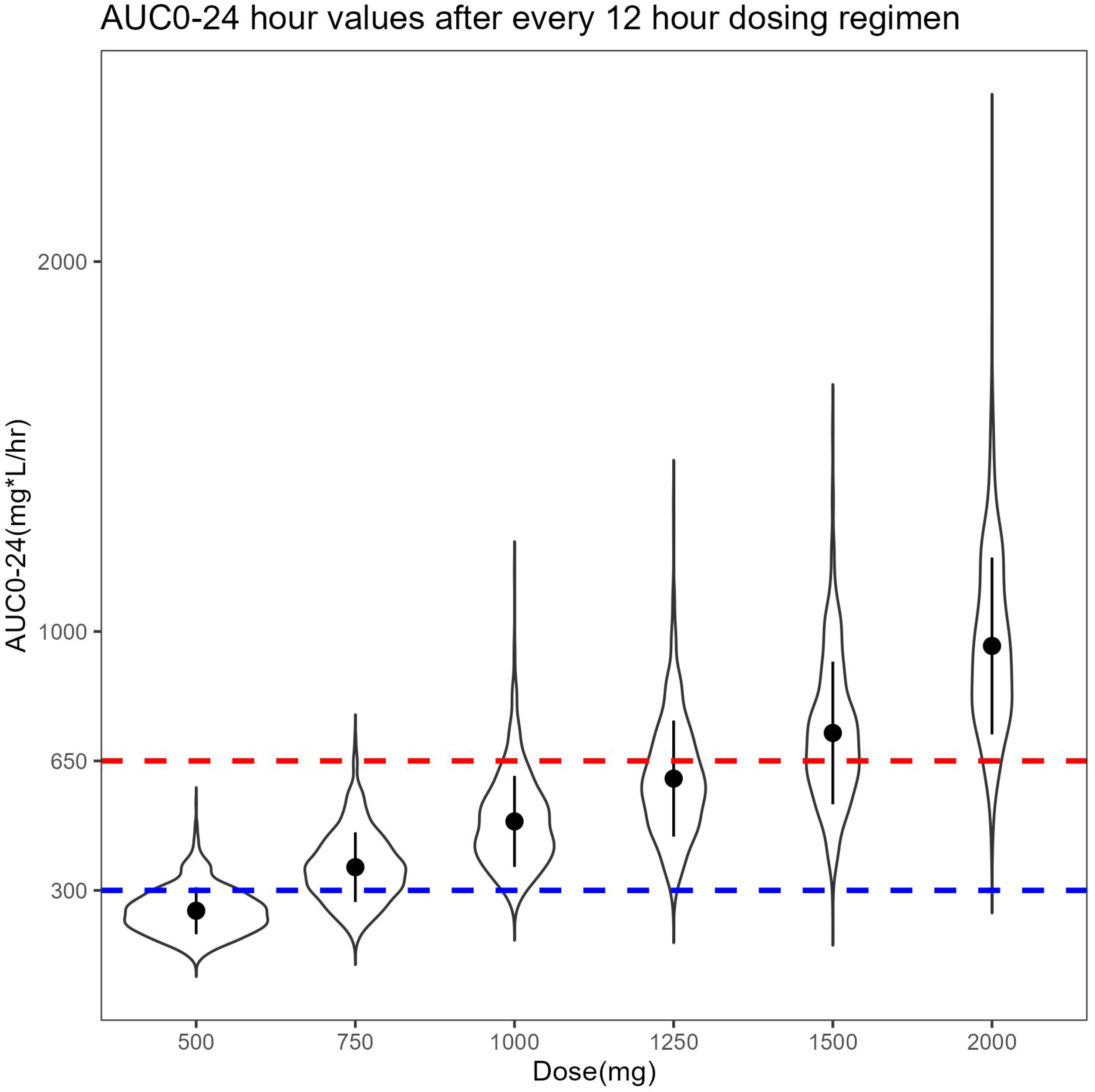
Violin plots of AUC_0-24_ versus various doses administered at every 12 hour interval. The black circle represents the median AUC_0-24_ and the black line represents the interquartile range. The white shaded area illustrates the kernel probability density.

## DISCUSSION

The American Thoracic Society Guidelines on Antibiotic Management of Lung Infections recommend IV vancomycin as the first line treatment for pulmonary exacerbations due to MRSA in PwCF (2). Even though, the pharmacokinetics of vancomycin is well studied in other populations (26-28), there are minimal data on PK of vancomycin in PwCF. Furthermore, the dosing of vancomycin is not consistent across multiple institutions across the United States and is dosed at a wide range between 500 to 2000 mg (6). In this study, we characterized the PK of vancomycin in PwCF with MRSA using a population pharmacokinetic approach based on Bayesian forecasting and applied the model to predict the PTA of an efficacy target of AUC/MIC ≥ 400 mg*h/L.

The PK of vancomycin in adult PwCF was reported in two studies (16, 17) and one study reported the PoPPK of vancomycin in pediatric population (29). Pleasants et al., 1996 evaluated the PK of vancomycin in PwCF after single-dose administration (15 mg/kg) and the PK samples were collected for only up to 24 h (16). In contrast, we evaluated the PK of vancomycin in PwCF across a period of 10 to 14 days after multiple-dose administration of vancomycin and included a broad range of dosing regimens (500 – 2000 mg q8h & q12h). In our study, vancomycin levels were at steady-state which is more representative of a clinical situation compared with the single-dose PK study reported by Pleasants et al. More recently, our group reported the PK of vancomycin in PwCF after intermittent dosing of vancomycin using the Sawchuk-Zaske method via two-point therapeutic drug monitoring (17). However, in this current study we used a more relevant Bayesian estimation approach to calculate the PK parameters. The pediatric PoPPK study by Stockmann et al., the median (range) age of the subjects 13.9 (8-17) years compared to 27 (19-60) years in our study indicating that both populations are significantly different from each other (29). Furthermore, in our study we applied our PoPPK model to evaluate the PTA of vancomycin for efficacy target of AUC_0-24_/MIC ≥ 400 and safety target of AUC_0-24_ < 650 after intermittent dosing, which was not addressed in all three reported studies in vancomycin PK in PwCF.

The single-dose PK study of vancomycin in PwCF reported a mean V_d_ of 40.6 L/70 kg and a mean CL of 5.7 L/hr/70 kg (16). Both these values are slightly higher than the values (V_d_ – 31.5 L/70 kg, CL – 5.5 L/hr/70 kg) reported in this study. The V_d_ calculated using the Sawchuk-Zaske method in recent study was 37.8 L/70 kg which is higher than the V_d_ (31.5 L/70 Kg) reported in this study (17). However, the CL was 5.1 L/hr/Kg which is quite similar to the CL (5.5 L/hr/70 kg) reported in this study. Nivia et al., 2022 in a scoping review reported the mean values of the PK parameters of vancomycin calculated using the PoPPK approach in non-critically ill adults (30). The PoPPK data was obtained from 17 studies and the median values of the estimated CL and V_d_ were 3.80 (interquartile range 2.84 to 5.42) L/h/70 kg) and 57.7 (interquartile range 46.9 to 65.7) L/70 kg. As expected, the PK values of vancomycin in PwCF from our study are different from the PK in non-critically ill adults with CL being higher (5.5 versus 3.8 L/hr/70 kg) and Vd being lower (31.5 versus 57.7 L/70 kg) (30). The difference in PK may be attributed to the pathophysiological changes in PwCF and further studies are warranted in this area.

Despite the fact that many PopPK models of vancomycin are described using two-compartment structural PK model (31, 32), controversy still exists on whether a one-or two-compartmental model is more appropriate to describe vancomycin PK from data obtained from routine therapeutic drug monitoring where data is sparse. In one study, the vancomycin PopPK model performance was compared between one- and two-compartment models using a Bayesian approach, and it was reported that the use of a two-compartment model resulted in statistically less bias and more precise predictions of vancomycin peak concentrations when either population parameters or non-steady-state concentrations were used for future predictions. However, they also concluded that no difference in model performance was observed when steady-state concentrations were used to predict future steady-state concentrations (33). Another study by Wu et al., compared the one-, two-, or three-compartment model performance of vancomycin, and suggested that for sparse data such as data from therapeutic drug monitoring (TDM) samples, one-compartment model is preferable (34). Similar to these reports, our data showed a better fit with a one-compartment model when compared with a two-compartment model. Our data is in agreement with Stockmann et al.’s report on vancomycin PK in pediatric PwCF, wherein, a one-compartment model was used to describe the TDM data of vancomycin in pediatric patients with PwCF (29).

In the covariate analysis, we evaluated the effect of co-administration of inhaled tobramycin and aztreonam on PK of vancomycin in PwCF. Both tobramycin and aztreonam are primarily excreted by kidneys after inhalation administration and may influence the PK of vancomycin (35). However, results showed that coadministration of inhaled tobramycin and aztreonam did not have significant effect on vancomycin PK. This is expected as the systemic absorption of both tobramycin and aztreonam were reported be very minimal and therefore may not have been reached at significant levels in plasma to alter the PK of vancomycin in PwCF (36, 37). Furthermore, in majority of health institutions, inhaled antibiotics are held off during the i.v antibiotic treatment in PwCF with PEx due to risk of nephrotoxicity. Our covariate analysis also showed that creatinine clearance did not significantly influence the PK of vancomycin which is an unexpected finding as the vancomycin is predominantly excreted unchanged by kidneys. The unexpected finding may be due to the short time period of 10-14 days of sample collection where kidney function of PwCF is not significantly altered and therefore not affecting vancomycin renal clearance.

The limitations of our study include use of data collected during routine TDM limiting the number of vancomycin concentrations measured for each patient, the short-time frame of sample collection, and not including the markers for efficacy such as MIC of vancomycin against MRSA in the patients included in the study and/or forced expiratory volume in 1 second (FEV1). Future studies should include prospective PK evaluation with more samples per patients for an extended time-period and include efficacy and toxicity data to correlate with the PK data and confirm the results of the model simulations.

In conclusion, vancomycin PK were adequately described by one-compartment model with first-order elimination in adult PwCF. The PK parameters V_d_ and CL in PwCF were not significantly different from non-critically ill patients. The Monte Carlo simulations showed that majority of the vancomycin dosing regimens currently used to treat pulmonary exacerbations in PwCF do not achieve the safety target and further studies must be conducted to investigate this issue. Future pharmacodynamic studies are needed to establish markers of efficacy and safety can be used to develop an optimal vancomycin dosing regimen to treat acute pulmonary exacerbations in adult PwCF.

## ACKNOWLDGEMENTS

We would like to thank Enhanced Pharmacodynamics LLC for providing an academic license of Finch Studio™ software.

